# Using computer vision to quantify facial expressions of children with autism during naturalistic social interactions

**DOI:** 10.1101/2025.05.14.653948

**Authors:** Liora Manelis-Baram, Tal Barami, Michal Ilan, Gal Meiri, Idan Menashe, Elizabeth Soskin, Carmel Sofer, Ilan Dinstein

**Author notes:** Corresponding Author : Liora Manelis-Baram.

## Abstract

**Background:** Difficulties with non-verbal communication, including atypical use of facial expressions, are a core feature of autism. Quantifying atypical use of facial expressions during naturalistic social interactions in a reliable, objective, and direct manner is difficult, but potentially possible with facial analysis computer vision algorithms that identify facial expressions in video recordings.

**Methods:** We analyzed >5 million video frames from 100 verbal children, 2-7-years-old (72 with autism and 28 controls), who were recorded during a ∼45-minute ADOS-2 assessment using modules 2 or 3, where they interacted with a clinician. Three different facial analysis algorithms (iMotions, FaceReader, and Py-Feat) were used to identify the presence of six facial expressions (anger, fear, sadness, surprise, disgust, and happiness) in each video frame. We then compared results across algorithms and across autism and control groups using ANCOVA analyses while controlling for the age and sex of participating children.

**Results:** There were significant differences in the performance of the three facial analysis algorithms including differences in the proportion of frames identified as containing a face and frames classified as containing each of the examined facial expressions. Nevertheless, analyses across all three algorithms demonstrated that there were no significant differences in the quantity of facial expressions produced by children with autism and controls.

**Limitations:** Our study was limited to verbal children with autism who completed ADOS-2 assessments using modules 2 and 3 and were able to sit in a stable manner while facing a wall mounted camera. In addition, this study focused on comparing the quantity of facial expressions across groups rather than their quality, timing, or social context.

**Conclusions:** Commonly used automated facial analysis algorithms exhibit large variability in their output when identifying facial expressions of young children during naturalistic social interactions. Nonetheless, all three algorithms did not identify differences in the quantity of facial expressions across groups, suggesting that atypical production of facial expressions in verbal children with autism is likely related to their quality, timing, and social context rather than their quantity during natural social interaction.

## Background

Facial expressions play a central role in non-verbal communication, conveying states, emotions, and intentions that are essential for effective social interaction [1,2]. Difficulties in non-verbal communication are a core symptom of autism [3], which can include the production of facial expressions that appear exaggerated, awkward, or flat and using facial expressions in different ways that impede social communication [4]. These difficulties may be associated with Alexithymia (i.e., difficulties identifying one’s own emotions), which appears in ∼50% of autistic individuals [5,6]. Despite the central role of facial expressions in social communication, relatively few studies have attempted to study them quantitatively in individuals with autism.

Previous studies have mostly used manual ratings or annotations to quantify differences in facial expressions across autism and control groups. For example, in some studies autistic and control individuals were explicitly instructed to pose or imitate specific facial expressions while they were recorded with video. When neurotypical [7,8,9] or autistic [10] individuals were asked to rate the recorded facial expressions, they reported that facial expressions produced by individuals with autism were similar in accuracy, but were more ambiguous, awkward, and atypical than those of controls.

Additional studies used manual coding schemes to quantify the production of spontaneous facial expressions in individuals with autism and controls. For example, two studies filmed children with autism and controls during short 8-minute social interactions with an adult clinician as they administered the Early Social-Communication Scales (ESCS) assessment [11]. The first study, using the Maximally Discriminative Movement Analysis coding system, reported fewer positive-affect expressions in children with autism compared to controls [12], while the second study, using the Affex coding system, reported no significant differences in expressions between groups [13].

A meta-analysis of facial expression studies that used a variety of manual techniques reported that individuals with autism exhibit fewer, shorter, less accurate, and more awkward facial expressions than controls [4]. Note that the majority of studies examined in this meta-analysis reported results from samples of <20 children in each group using analysis of video recordings that were extremely short. Since facial expressions are likely to vary across children and over time, extending these studies to longer recordings from larger cohorts is critical for establishing generalizable conclusions. This will require utilization of automated techniques with facial analysis algorithms.

Over the last decade multiple computer vision algorithms with the ability to identify facial expressions in video recordings have been released [14] including iMotions FACET [15], iMotions AFFDEX [16], OpenFace [17], FaceReader [18], and Py-Feat [19]. Several studies have applied these algorithms to analyze videos of individuals with autism who were explicitly asked to pose or imitate facial expressions. One study using the iMotions FACET algorithm reported that posed facial expressions involved weaker muscle contractions (i.e., known as action units - AUs) in the autism relative to the control group particularly when expressing happiness [20]. However, other studies found no significant difference in the intensity of action unit activations across groups in any of several posed/imitated facial expressions when analyzed with OpenFace [17,21]. When applying the iMotions FACET algorithm to video recordings of participants watching movies, one study reported that individuals with autism exhibited more neutral facial expressions than controls [5], while another did not find any differences across groups [22]. A third study using a custom-built algorithm also reported more neutral facial expressions in children with autism than controls as they watched a series of movie clips [23].

Only three studies to date have used automated algorithms to examine spontaneous facial expressions during naturalistic social interactions, a condition where individuals with autism are expected to exhibit the largest difficulties. The first used OpenFace to analyze video recordings of 10-minute conversations between adolescents and their mothers or a female research assistant [24]. In comparison to controls, adolescents with autism exhibited significantly fewer facial action unit activations when smiling and poor facial synchronization with the research assistant, but not with their mother. Another study examined videos of a 7-minute dialog between adults and an actress, also using OpenFace [25]. This study reported that adults with autism exhibited less frequent mimicry (i.e., synchronization) than controls, less frequent activation of smiling action units in parts of the dialog intended to evoke positive emotions, and more frequent activation of disgust action units in parts of the dialog intended to evoke negative emotions. Finally, a third study used FaceReader to quantify facial expressions of adults in ∼1-minute segments of video recorded during the cartoon task of the ADOS-2, module 4 assessment. They reported more neutral and fewer happy facial expressions in the autism group relative to controls [26].

Note that these studies examined short video segments with different facial analysis algorithms and reported mixed results regarding potential differences in the quantity, quality, and synchronization of facial expressions produced by individuals with autism. Moreover, these studies were performed with adolescents and adults and only one study used video recordings from ADOS-2 assessments, which are commonly available in many clinical and research settings and offer a standardized semi-structured context for evaluating social communication difficulties.

The current study had several goals. First, we wanted to examine facial expressions in young children with autism (2-6-years-old) rather than adolescents/adults given that the production of facial expressions may change as individuals develop and potentially compensate for early difficulties. Second, we wanted to assess the reliability of quantified facial expressions by comparing results across three commonly used algorithms: iMotions, FaceReader, and Py-Feat. Third, we wanted to quantify facial expressions in considerably longer video recordings (∼45 minutes) than those used in previous studies (1-10 minutes) given that social interactions are dynamic, and the production of facial expressions may vary from one minute to another. Fourth, we wanted to include a relatively large number of children with autism given that individuals may vary considerably in their production of facial expressions. Fifth, we wanted to quantify facial expressions within a structured context that could be easily replicated by other labs and clinics. To achieve these goals, we analyzed full recordings of ADOS-2 assessments (∼45 minutes) and assessed the agreement across the three algorithms in identifying and quantifying facial expressions. Most importantly, we compared the proportion of facial expressions produced by autistic and control children to determine whether there were consistent significant differences in the quantity of facial expressions across groups.

## Methods

### Participants

We extracted video recordings of 100 children from the National Autism Database of Israel (NADI) managed by the Azrieli National Centre for Autism and Neurodevelopment Research (ANCAN). ANCAN is a collaborative project between Ben Gurion University (BGU) and eight clinical sites throughout Israel [27]. All recordings analyzed in this study were performed at the Soroka University Medical Center (SUMC). NADI contains video recordings of clinical assessments, along with various other behavioral and clinical measures from a growing cohort of children with autism in Israel [28]. All children were recruited between 2018 and 2024 and their parents completed informed consent. This study was approved by the Helsinki committee of SUMC and the IRB committee of BGU.

The sample included 72 children with autism (17 girls), 3.16-6.91 years old, and 28 typically developing control children (8 girls), 2.58-6.66 years old (Table 1). All children completed the Autism Diagnostic Observation Schedule, Second Edition (ADOS-2) using module 2 or 3 [29]. All children with autism and none of the control children exceeded the ADOS-2 cutoff for autism and met the *Diagnostic and Statistical Manual of Mental Disorders*, Fifth Edition (DSM-5, [3]) criteria for autism, as determined by both a physician (child psychiatrist or pediatric neurologist) and a developmental psychologist. In addition, 86 of the children completed a developmental or cognitive assessment as appropriate for their age. Two children completed the Bayley scales of infant and toddler development, 3^rd^ edition [30], 46 children completed the Mullen scales of early learning [31], and 38 completed the Wechsler preschool and primary scale of intelligence (*WPPSI-III,* [32]).

**Table 1.** Descriptive Statistics of the participating children including their age, sex, ADOS-2 scores and Developmental\Cognitive score, per group, and comparison across groups.

|  | Autism (n=72) | Control (n=28) | Statistics |
| --- | --- | --- | --- |
|  | Mean (std) | Mean (std) |  |
| **Age (years)** | 56.03 (12.35) | 46.11 (13.07) | t(98) = 3.55, p<.001 |
| **Sex (girls)** | n=17, 23.61% | n=8, 28.57% | $\chi^2(1) = .08$, p= .76 |
| **ADOS-2 scores\*** |  |  |  |
| Total Calibrated Severity Score (CSS) | 6.18 (1.68) | 1.5 (0.69) | t(98) = 8.18, p<.001 |
| Social Affect (SA) CSS | 5.49 (2.07) | 2.07 (1.21) | t(99) = 8.18, p<.001 |
| Restricted Repetitive Behaviors (RRB) CSS | 7.78 (1.7) | 1.64 (1.62) | t(98) = 16.44, p<.001 |
| **Developmental\Cognitive scores\*\*** | 85.41 (17.38) | 111.11 (12.63) | t(84) = -6.88, p<.001 |

\*\* Developmental\Cognitive scores were available for 86 children, 59 autism and 27 controls. std=standard deviation

### Video recordings

All children were recorded during an ADOS-2 assessment, a semi structured standardized diagnostic test for identifying autism. A clinician with research reliability selected the appropriate ADOS-2 module based on the child’s age and level of expressive language. Recorded ADOS-2 sessions were 32.31-75.9 minutes long (M=53.19, SD=8.76) and did not differ across autism and control groups (autism: M=53.6, SD=9.32, control: M=52.12, SD=6.78, t(98)=0.75, p=0.45). As noted above, we intentionally selected recordings of children who completed ADOS-2 modules 2 and 3 because they require the child to sit at a table, making their face clearly visible to a nearby camera. The ADOS-2 video recordings were performed in five different assessment rooms installed with similar camera systems at SUMC.

### Facial expression analysis

We processed the video recordings using three facial analysis software packages that extract a similar set of facial expressions from video data:

#### iMotions

Commercially available software package that utilizes the AFFDEX algorithm [16] to detect a face and the presence of 6 facial expressions on each video frame (joy, anger, sadness, disgust, surprise, and fear). It returns a probabilistic score (0-100) representing the degree of confidence that each facial expression was present in the frame. Emotion scores were rescaled to a range of 0-1 to fit the range of the other algorithms.

#### FaceReader (Noldus Inc.)

Commercially available software package that utilizes a proprietary computer vision algorithm to detect a face and the presence of 7 facial expressions (happy, angry, sad, disgusted, surprised, scared, and neutral). FaceReader fits a mesh with ∼500 vertexes to the face and computes a probabilistic score (0-1) that a certain facial expression is present in each frame.

#### Py-Feat

Free, open-source python toolkit that integrates multiple algorithms for detection of faces, facial landmarks, facial action units, and facial expressions [19, 33]. We used the img2pose algorithm for face detection [34], which yields a confidence score (0-1) for the detection of a face per frame. We used the Resmasknet algorithm [35] for detecting 7 facial expressions (happiness, anger, disgust, fear, sadness, surprise, and neutral). Unlike the two commercial algorithms, Resmasknet scores each of the 7 facial expressions with a probabilistic score (0-1) that represents their likelihood in relative terms such that the sum of their scores equals one.

### Preprocessing

Since there were typically multiple individuals present in the assessment room and captured in the video recordings (i.e., child, parent, and clinician), we manually placed a bounding box around the child in each movie. Consequently, all data outside the bounding box was excluded from analysis. While each software required a separate definition of the bounding box, we used room landmarks to ensure the bounding box was placed in the same location across all three.

Each algorithm detected a face within the specified bounding box in some, but not all the frames (e.g., when the child left the bounding box or face landmarks were not visible). To ensure that the analyzed data included videos with reliable and continuous face detection, we excluded frames according to the following criteria. First we extracted the pitch, roll, and yaw of the child’s head per frame from each algorithm and excluded all frames with values above 75 degrees relative to the camera in any direction (i.e., child was facing away from the camera). Second, we excluded isolated video segments, shorter than 25 frames (approximately 1 second), that were preceded and followed by frames without valid face detection. Third, the img2pose algorithm in Py-Feat, unlike the other two algorithms, also reported a confidence score for face detection per frame and we excluded frames with a face score below 0.9.

Finally, we applied a low-pass Gaussian filter with a width of 11.3 frames (Approximately 0.5 of a second at half-height) to smooth the time-course of each facial expression, thereby minimizing rapid changes in facial expressions that are likely to result from measurement noise.

### Facial analysis measures

All data analysis was performed with custom written code in Python. First, we computed the number of valid frames where a face was detected within the child’s predefined bounding box per video. We compared both the absolute number of valid frames (in minutes) and their proportion (i.e., valid face frames divided by total number of frames in the video). Second, we computed the number of frames where a given facial expression exceeded a value of 0.5 (same threshold for all algorithms). This analysis was performed separately for each of six facial expressions: anger, fear, sadness, surprise, disgust, and happiness. For each facial expression, we computed its proportion relative to the total number of valid face frames per video. These proportions were compared across algorithms and across participant groups to ensure that the results were not influenced by differences in the length of recorded ADOS assessments or the proportion of valid face frames. Finally, we extracted frame-by-frame happiness time-courses from each algorithm, which contained scaled values of 0-1 representing the confidence of the algorithm that a happiness facial expression was exhibited by the child on a given video frame. We then calculated pair-wise correlations across algorithms per video, enabling us to assess their frame-by-frame agreement in identifying happiness.

### Statistical analysis

All statistical analyses were performed using custom written code in Python. Student t-tests were used to assess age and behavioral score differences across autism and control groups. A Chi square test was used to compare the proportion of males and females across groups. Analysis of Covariance (ANCOVA) tests were used to compare facial analysis results across algorithms and participant groups while using Tukey’s Honestly Significant Difference (HSD) tests for post hoc analyses of between-group differences. Pearson correlation coefficients were computed to assess pair-wise algorithm agreement in quantifying happiness across children. Pearson correlations were also used to assess the similarity of facial expression time-courses across algorithms. All statistical tests were performed with a significance level set to α = 0.05.

## Results

We first compared the ability of the three algorithms to detect the child’s face on individual frames of each movie. ANCOVA analyses with age, sex, and diagnosis as covariates revealed that face detection differed significantly across the three algorithms when comparing either the absolute number of valid face frames or their proportion relative to video length (Table 2). Tukey’s HSD Post hoc test demonstrated that the absolute number of valid face frames detected by iMotions (M = 21.76, SD = 9.35) was significantly lower compared to FaceReader (M = 31.61, SD = 9.69, p = 0.001) and Py-Feat (M = 31 19, SD = 10.25, p = 0.001). Similarly, the proportion of valid face frames detected by iMotions (M = 0.43, SD = 0.16) was significantly lower compared to FaceReader (M = 0.63, SD = 0.16, p < 0.001) and Py-Feat (M = 0.62, SD = 0.17, p < 0.001). There were no significant differences between FaceReader and Py-Feat in either measure.

**Table 2.**
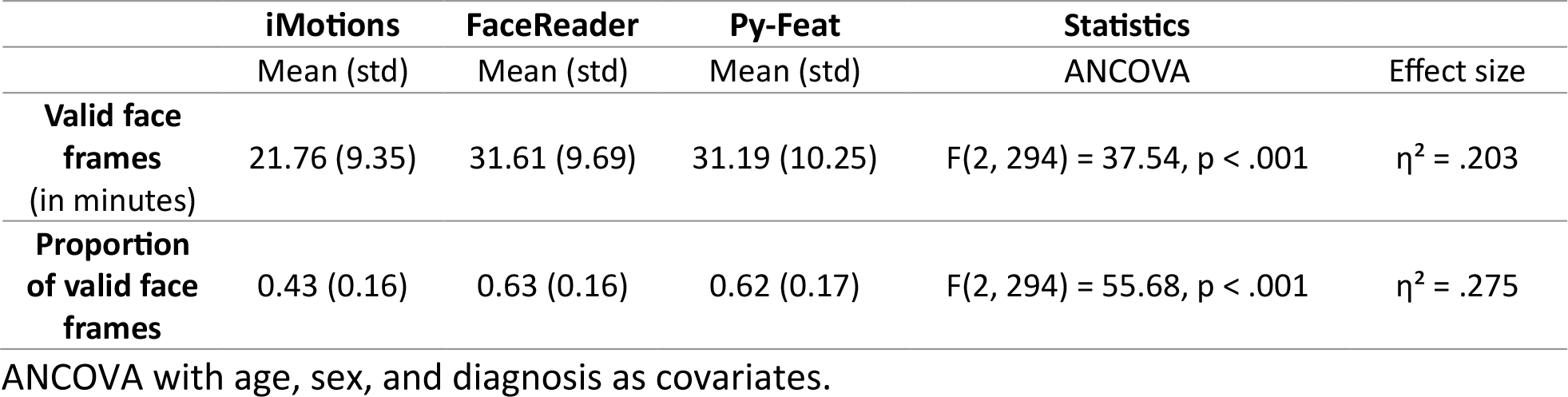
Comparison of the number and proportion of valid face frames across algorithms.

|  | iMotions | FaceReader | Py-Feat | Statistics |  |
| --- | --- | --- | --- | --- | --- |
|  | Mean (std) | Mean (std) | Mean (std) | ANCOVA | Effect size |
| **Valid face frames**<br>(in minutes) | 21.76 (9.35) | 31.61 (9.69) | 31.19 (10.25) | $F(2, 294) = 37.54$, $p < .001$ | $\eta^2 = .203$ |
| **Proportion of valid face frames** | 0.43 (0.16) | 0.63 (0.16) | 0.62 (0.17) | $F(2, 294) = 55.68$, $p < .001$ | $\eta^2 = .275$ |

ANCOVA with age, sex, and diagnosis as covariates.

The absolute number and proportion of valid face frames was larger in controls than children with autism (absolute number: F(1, 294) = 37.43, p < 0.001, η^2^ = 0.113, proportion: F(1, 294) = 59.19, p < 0.001, η^2^ = 0.168) and larger in older versus younger children (absolute number: F(1, 294) = 27.33, p < 0.001, η^2^ = 0.085, proportion: F(1, 294) = 21.46, p < 0.001, η^2^ = 0.068). There were no significant differences between boys and girls. This indicated that diagnosis and age, but not sex, had an impact on successful face detection by the algorithms.

### Facial expression analyses

Next, we compared the proportion of frames with facial expressions of anger, fear, happiness, sadness, surprise, and disgust across the algorithms. ANCOVA analyses revealed significant differences across algorithms for all facial expressions, but not for age, sex, or diagnosis covariates (Table 3). These analyses revealed that happiness was the most common facial expression consistently identified by all three algorithms.

**Table 3.** Proportion of frames with identified facial expressions per algorithm.

|  | iMotions | FaceReader | Py-Feat | Statistics |  |
| --- | --- | --- | --- | --- | --- |
|  | Mean (std) | Mean (std) | Mean (std) | ANCOVA | Effect size |
| **Happiness** | 4.41 (4.32) | 11.35 (6.99) | 18.74 (12.50) | $F(2, 294) = 68.75, p < .001$ | $\eta^2 = .319$ |
| **Anger** | 0.15 (0.16) | 0.49 (0.38) | 1.03 (1.05) | $F(2, 294) = 46.74, p < .001$ | $\eta^2 = .241$ |
| **Disgust** | 0.32 (0.49) | 0.91 (0.73) | 1.07 (2.36) | $F(2, 294) = 7.37, p < .001$ | $\eta^2 = .048$ |
| **Fear** | 0.09 (0.10) | 0.48 (0.39) | 5.73 (6.41) | $F(2, 294) = 73.20, p < .001$ | $\eta^2 = .332$ |
| **Sadness** | 0.21 (0.95) | 4.49 (4.14) | 22.59 (16.89) | $F(2, 294) = 138.46, p < .001$ | $\eta^2 = .485$ |
| **Surprise** | 0.91 (1.42) | 1.41 (1.53) | 5.02 (7.04) | $F(2, 294) = 28.08, p < .001$ | $\eta^2 = .160$ |

Proportion of frames exceeding a confidence threshold of 0.5 (see methods). ANCOVA with covariates for age, sex, and diagnosis.

To assess algorithm agreement per child, we performed follow-up correlation analyses with the happiness facial expression given its prominence. We extracted the proportion of happiness frames per child, per algorithm, and computed both Concordance and Pearson correlation coefficients across algorithm pairs, separately for the autism and control groups (Figure 1).

**Figure 1.**
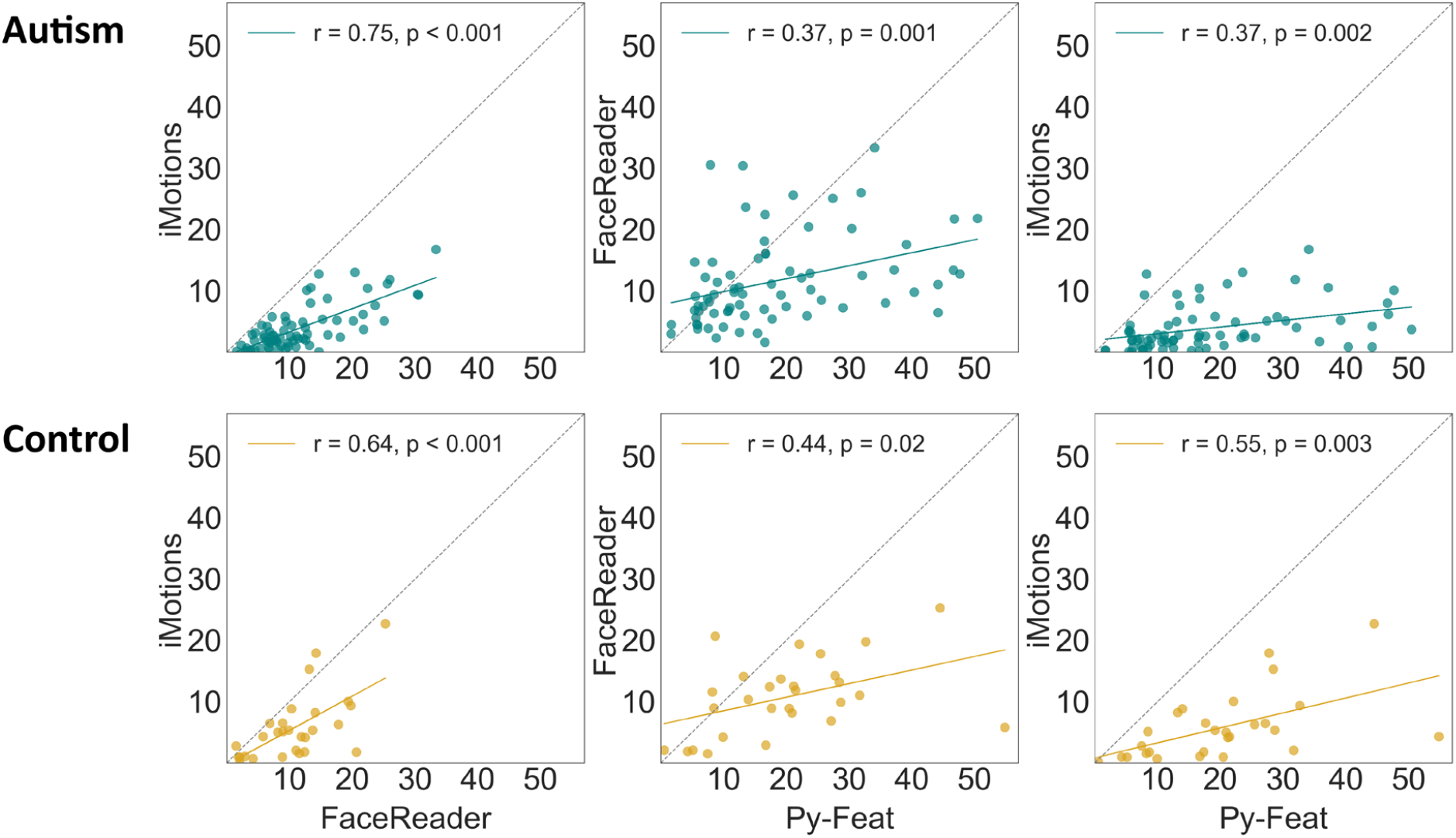
Scatter plots demonstrating the correlation across algorithm pairs in the proportion of happiness identified per child. Left column: iMotions and FaceReader. Middle column: FaceReader and Py-Feat. Right column: iMotions and Py-Feat. Each point represents one child. Green: autism. Yellow: controls. Dotted line: unity line. Solid line: least squares linear fit.

Pearson correlations between FaceReader and iMotions were relatively high (autism: r=0.75, p < 0.001, control: r=0.64, p<0.001) and correlations of FaceReader with Pyf-Fat (autism: r=0.37, p=0.001, control: r=0.44, p=0.02) or iMotions with Py-Feat (autism: r=0.37, p=0.002, control: r=0.55, p=0.003) were lower. While all Pearson correlations were significant, most effect sizes were in the low to medium range, suggesting relatively weak agreement across algorithms.

### Happiness time courses

To examine agreement across algorithms further, we extracted frame-by-frame time-courses of happiness per child, per algorithm. Each time-course contained scaled values of 0-1 representing the confidence of the algorithm that a happiness facial expression was exhibited by the child on a given video frame. We then computed the Pearson correlation across time-courses of algorithm pairs to assess their temporal agreement. These analyses revealed heterogeneous results across children/recordings (Figure 2). For some children the three algorithms exhibited strong correlations that were 3-4 times stronger than others. An ANCOVA analysis revealed significant differences in the correlation values (i.e., agreement) of specific algorithm pairs (F(2, 294) = 24.91, p < 0.001, η^2^ = 0.145) with Tukey’s HSD tests indicating that FaceReader-iMotions (p<0.001)) and FaceReader–Py-Feat (p<0.001) exhibited significantly higher time-course correlations compared to iMotions-Py-Feat. The diagnosis covariate was also significant (F(1,98)=4.56,p=0.036) such that agreement across algorithms was higher for controls. Age and sex covariates were not significant covariates (p>0.05).

**Figure 2.**
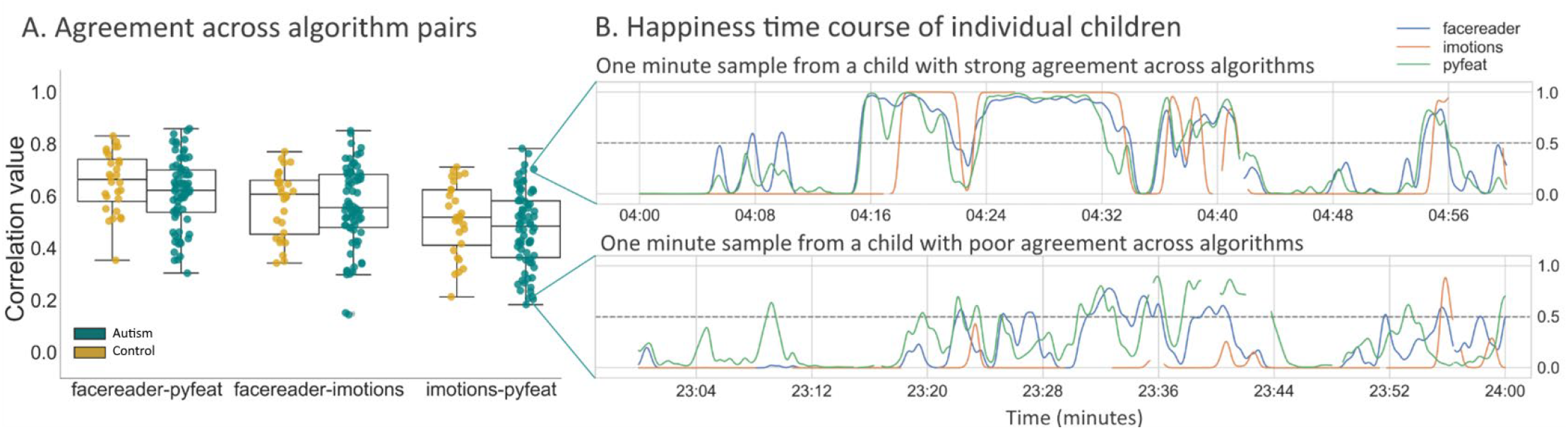
Agreement across algorithms in identifying happiness in frame-by-frame movie time-courses. **A.** Scatter plot with time-course correlation values per child, demonstrating the degree of agreement across pairs of algorithms for individual children. Yellow: controls. Green: autism. **B.** One-minute sample of happiness time-course extracted from a child with strong agreement across iMotions and Py-Feat algorithms. **C.** One-minute sample from a child with poor agreement. Blue: FaceReader. Orange: iMotions. Green: Py-Feat.

### No differences in the quantity of facial expressions between autism and control children

We performed ANCOVA analyses to compare the proportion of frames with each of the 6 facial expressions across participant groups while controlling for age, sex and proportion of valid face frames (i.e., frames where a face was detected). These analyses revealed that there were no significant differences in the proportion of facial expressions across groups regardless of algorithm (Table 4).

**Table 4.**
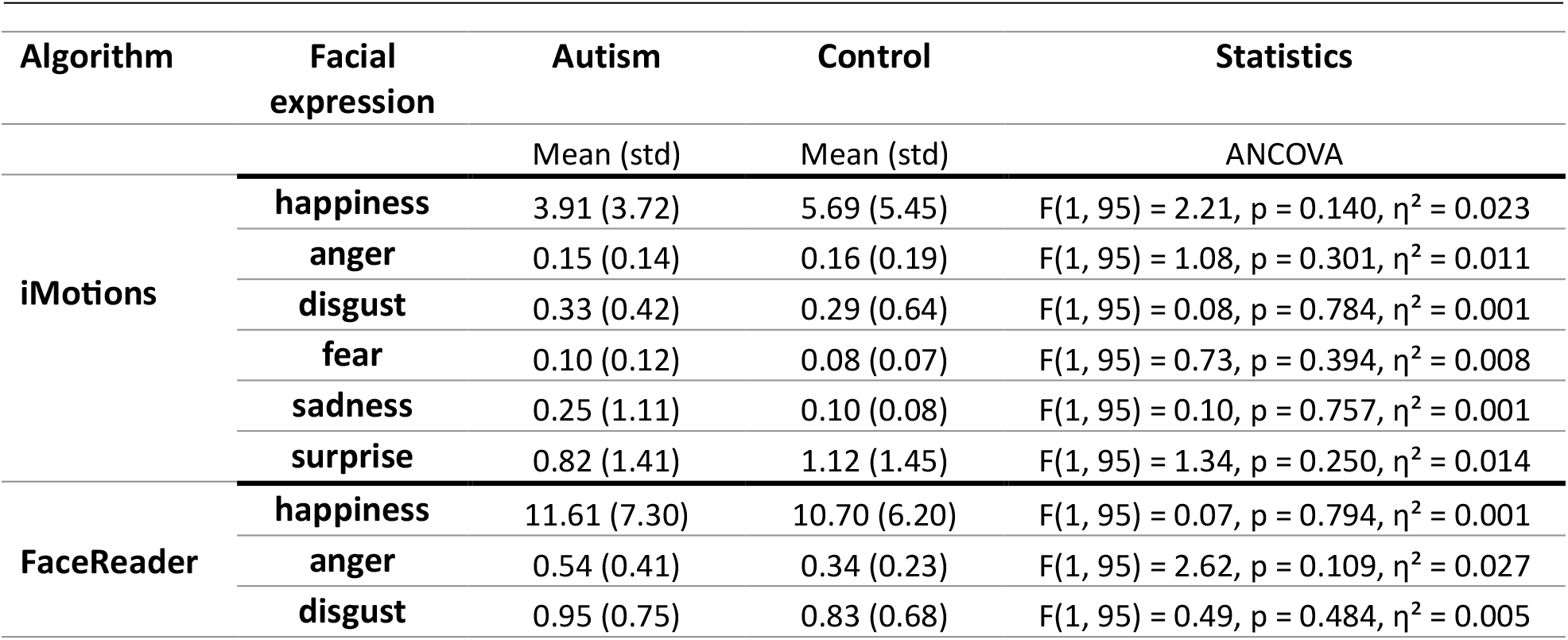
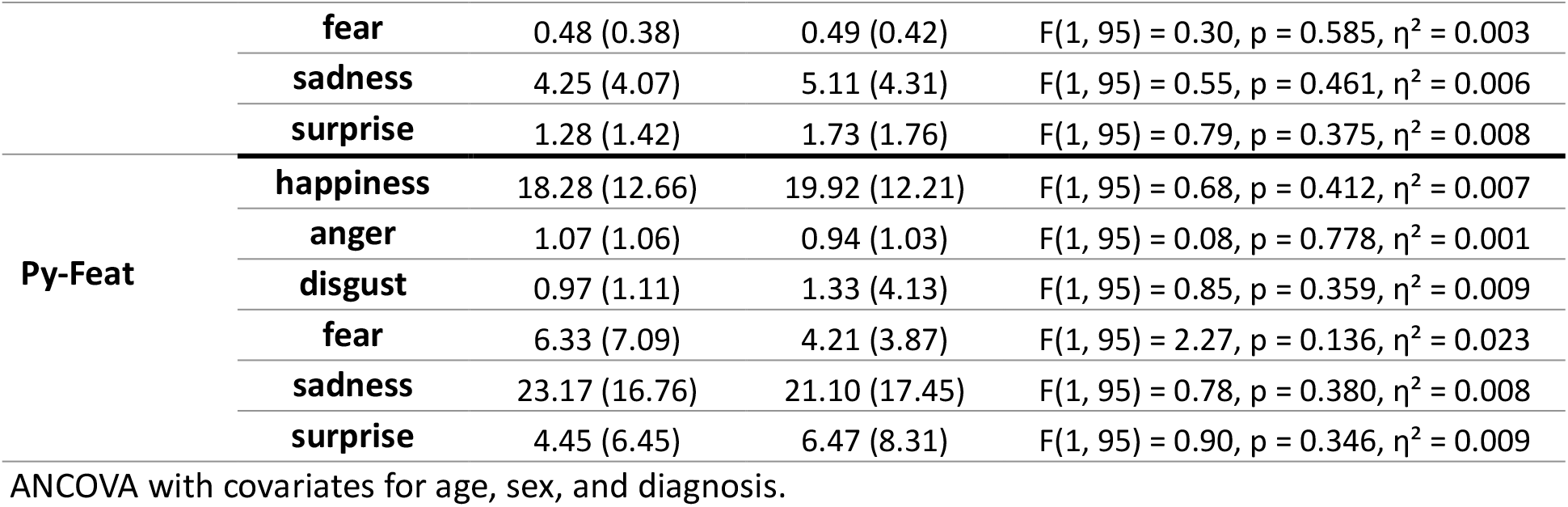
Comparison of the proportion of frames with facial expression across autism and control groups. Analyses were performed separately for each of the three algorithms.

### Relationship with autism severity

In a final analysis we examined whether overall production of facial expressions was related to the severity of core autism symptoms as estimated by ADOS-2 CSS scores. We calculated the proportion of frames with any facial expression per subject and computed Pearson correlation with total ADOS-2 CSS scores separately for each diagnostic group (Figure 3). There were no significant correlations when performing this analysis with FaceReader (ASD: r=-0.15, p = 0.2, control: r=0.06, p=0.77), iMotions (ASD: r=-0.08, p=0.5, control: r=-0.15, p=0.46) or Py-Feat (ASD: r=0.11, p=0.36, control: r=-0.31, p=0.11). These findings suggest that the quantity of facial expressions was not consistently related to the severity of core autism symptoms as estimated by the ADOS-2.

**Figure 3.**
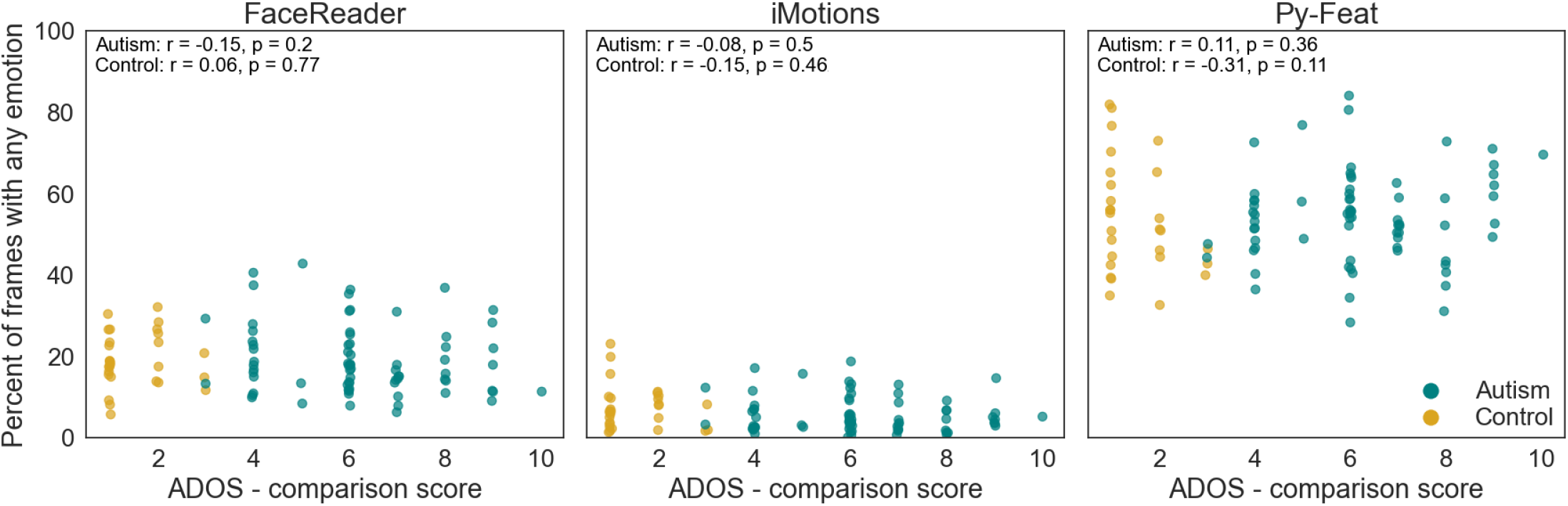
Relationship between the percent of frames with any emotion detected above a threshold of 0.5 (y-axis) and ADOS - comparison scores (x-axis) by algorithm (FaceReader, iMotions, and Py-Feat). Points represent data for individual children in the autism (green) and control (yellow) groups.

## Discussion

All children with autism exhibit difficulties in non-verbal social communication, by definition, since it is a core diagnostic feature. However, these difficulties can be manifested in many ways that may or may not include atypical production of facial expressions. Moreover, atypicalities may include differences in the quantity, quality (e.g., shape or temporal dynamics), timing, or social context of facial expressions.

The present study shows that both children with autism and controls exhibit large heterogeneity in the quantity of spontaneous facial expressions produced during a structured social interaction with a clinician (Figure 1). However, the quantity of facial expressions did not differ significantly between the two groups in any of the examined facial expressions, including happiness, anger, disgust, fear, sadness, or surprise (Table 4). These results were consistent across three different facial analysis algorithms (iMotions, FaceReader, and Py-Feat) and suggest that potential facial expression atypicalities in children with autism are not necessarily apparent in their quantity during a social interaction.

### Facial expression atypicalities in autism

Previous studies, using a variety of different techniques and experimental designs, have reported mixed results with some reporting no significant differences in the amount of facial expressions produces by individuals with autism relative to controls [13], as we also report here, and others reporting reduced frequency of smiling and more neutral expressions in the autism group [12,24,25,26]

There are multiple factors that may explain differences between previously reported results and those in our study. First, previous studies examined considerably shorter video recordings of social interaction (1-8 minutes) that may have yielded serendipitous findings by chance due to the limited video sample examined [25, 26]

24]. The duration of video samples in our study was much longer (53 minutes on average), providing a considerably larger opportunity to identify facial expressions than possible in previous studies. Since facial expression frequency may vary over time, extensive sampling is needed for establishing reliability. Indeed, future studies would benefit greatly from multi-day recordings per subject to establish test-retest reliability.

Another factor is the context of the video recording, which is likely to influence the amount and type of facial expressions exhibited by the children. Some of the studies that reported differences across groups used unique contexts involving specific scripted tasks or dialogues with a research assistant [25, 24]. Others utilized specialized assessments, such as the Early Social Communication Scales [12, 13), which require training and are not commonly administered in clinical or research settings. Our study utilized recordings of the ADOS-2 assessment which constrains the recordings to a reproducible context that is widely used and easy to replicate although it also requires training and maintaining research reliability.

A third factor that may explain differences between our study and previous ones is the age of the participants. Our participants were relatively young children (ages 2–8-years-old), in contrast to other studies that recruited adolescents [24] and adults [25,26]. The ability to produce facial expressions develops continuously throughout childhood and adolescence in the general population, with full maturity typically achieved in late adolescence [36,37]. Hence, differences across autism and control groups may vary with age as a function of facial expression maturity and the ability of the algorithms to accurately identify younger facial expressions.

### Differences across facial analysis algorithms

Advances in computer vision and machine learning techniques have led to the development of multiple automated algorithms for the identification of facial expressions [38]. While some algorithms may yield high accuracy (∼90%) when applied to video recordings of adults in controlled lab settings where the participant faces the camera, performance drops significantly (∼50%) when applied to “real world” scenarios [39]. Factors such as lighting, angle of the face relative to the camera, and partial occlusions (e.g., wearing glasses or a hat) are key challenges [14]. Since most algorithms were trained with images of adults performing specific facial expressions, which often include posed and exaggerated expressions [40,41), their ability to accurately identify facial expressions of children during naturalistic interactions may not be as high as often claimed by commercial vendors.

An important contribution of the current study was examining the reliability of automated facial expression analyses across different algorithms. We found significant systematic differences in the algorithms’ ability to detect faces and identify facial expressions. FaceReader and Py-Feat detected faces in 50% more frames than iMotions (Table 2), and Py-Feat identified emotions in three to four times as many frames as FaceReader and iMotions (Table 3 and Figure 3). These findings extend previous reports from studies with typically developing adults demonstrating variability in the output of different facial analysis algorithms [14]. Developing more reliable and accurate algorithms would be critical for studying the timing, synchrony, and quality of facial expressions given that such differences are likely embedded in the moment-by-moment dynamics of facial expressions (Figure 2). Addressing this challenge will require open-science repositories containing diverse, manually annotated videos of children in naturalistic social interactions. These data would serve as “ground truth” datasets that would enable training and testing of new and existing algorithms.

Our study emphasizes the poor reliability apparent across three leading algorithms used to analyze the same set of video recordings. Dramatic differences were apparent in the ability of the algorithms to identify the child’s face and facial expressions on individual frames (Tables 2 and 3), and there was poor agreement across algorithms when quantifying the total amount of smiles exhibited by individual children as well as their temporal time-courses, particularly in the autism group (Figures 1 and 2).

These results highlight an urgent need to develop new open-source facial analysis algorithms that are specifically trained to identify the facial expressions of young children filmed in naturalistic interactions. Achieving this goal will require the establishment of large video repositories with accurately annotated data that can be accessed freely by researchers (e.g., [42]).

### Limitations

Our study had several limitations. First, all participating children completed ADOS-2 assessments using modules 2 and 3, which require the child to sit at a table and engage in verbal interactions. As such, our findings are relevant only to verbal children with autism. Second, we focused our analyses on quantifying the presence of specific facial expressions, rather than measuring their quality, timing, or social context (e.g., directed towards another individual in a communicative manner or not). Third, while our sample size was one of the largest to date, it is still small for capturing the large heterogeneity clearly apparent across children with autism and controls. Indeed, the promise of automated facial analysis algorithms is to scale such analyses to thousands of participants with extensive multi-day video recordings per subject that would be necessary to establish test-retest reliability. Finally, our findings regarding poor reliability across algorithms suggest that these are early days for automated facial analysis studies of children with autism and that an effort to create large annotated video repositories of children with autism for further algorithm development is highly warranted.

## Conclusions

Our findings indicate that young children with autism produce similar quantities of spontaneous facial expressions during naturalistic social interactions compared to their typically developing peers. This suggests that difficulties in non-verbal social communication in verbal children with autism may be more related to the quality, timing, or contextual appropriateness of facial expressions rather than their overall frequency. Significant variability in the output of the three facial analysis algorithms highlights the current limitations of automated facial expression recognition, particularly when applied to young children in naturalistic settings. These results emphasize the need to develop more reliable open-source algorithms and large, annotated video repositories that can serve as ground truth datasets for training and testing. Such advancements are critical for enabling robust and scalable studies of facial expressions in autism, ultimately contributing to better diagnostic tools and interventions.

## Acknowledgments

We thank all the children and families who participated in this study. We also acknowledge the clinicians and research assistants for their assistance with data collection and ADOS-2 assessment administration.

## List of abbreviations

ASD: Autism Spectrum Disorder

ADOS-2: Autism Diagnostic Observation Schedule, Second Edition

## Declarations

### Ethics approval and consent to participate

This study was approved by the Helsinki committee of SUMC and the IRB committee of BGU. All parents completed informed consent.

## Consent for publication

Not applicable.

## Contributions

L.M.B., I.D. conceived and designed the study and wrote the main manuscript text. L.M.B. and M.I. performed the clinical assessments and data acquisition. L.M.B., T.B., and E.S. conducted the data processing and analyses. I.M., G.M. and C.S. critically revised the manuscript for important intellectual content. All authors reviewed and approved the final version of the manuscript.

## Availability of data and materials

Data and materials are available on request.

